# Autism-linked gene FoxP1 selectively regulates the cultural transmission of learned vocalizations

**DOI:** 10.1101/2020.03.14.992016

**Authors:** Francisco Garcia-Oscos, Therese Koch, Harshida Pancholi, Massimo Trusel, Vamsi Daliparthi, Fatma Ayhan, Marissa Co, Danyal H. Alam, Jennifer E. Holdway, Genevieve Konopka, Todd F. Roberts

**Author notes:** Equal contribution (Francisco Garcia-Oscos, Therese Koch, Harshida Pancholi, Massimo Trusel). Correspondence should be addressed to TFR.

## Abstract

Autism spectrum disorders (ASD) are characterized by impaired learning of culturally transmitted behaviors like social skills, speech, and language^1–3^. These behaviors are learned by copying parents and other social models during development, a two-stage process that involves forming memories of appropriate behaviors during social experiences and then using those memories to guide imitation. How ASD-linked genes impair these often-intertwined aspects of learning is not known, thereby limiting our understanding of the developmental progression of ASD and the targeting of therapeutic interventions. Here we show that these aspects of learning are dissociable and that the ASD-linked gene *FoxP1* selectively impairs learning from social experience, but not behavioral imitation. Haploinsufficiency of *FOXP1* in humans causes FOXP1 syndrome, a neurodevelopmental disorder typified by severe disruptions in speech and language development, and other ASD-associated symptoms^4,5^. We tested how knockdown of *FoxP1* (FP1-KD) affects the cultural transmission of vocal behaviors in zebra finches, a songbird that learns by memorizing and vocally copying the song of an adult ‘song-tutor’. We find that FP1-KD blocks song learning in juvenile birds by selectively impairing their ability to encode a memory during social experiences with a song-tutor. These learning deficits are linked to disruptions in experience-driven structural and functional plasticity. However, if birds are exposed to tutor-song prior to FP1-KD, their ability to imitate that song during development is unaffected. Thus, FP1-KD impairs cultural transmission of vocalizations by disrupting the ability to form appropriate vocal memories, yet spares the ability to use previously acquired memories to guide vocal learning. This indicates that learning from social experience may be particularly vulnerable in FOXP1 syndrome.

---

Humans and other animals learn many of their complex and socially oriented behaviors by imitating more experienced individuals in their environment. For example, development of spoken language is rooted in a child’s ability to imitate the speech patterns of their parent(s) and other adults ^6–8^. This cultural transmission of behavior is impaired in many neurodevelopmental disorders, most notably ASD^1–3^. However, how ASD risk genes impact behavioral imitation is still not known. We sought to examine this issue by testing the role of FoxP1 in the cultural transmission of song between adult and juvenile zebra finches (Fig. 1a-d, Extended Data Fig. 1).

**Figure 1.**
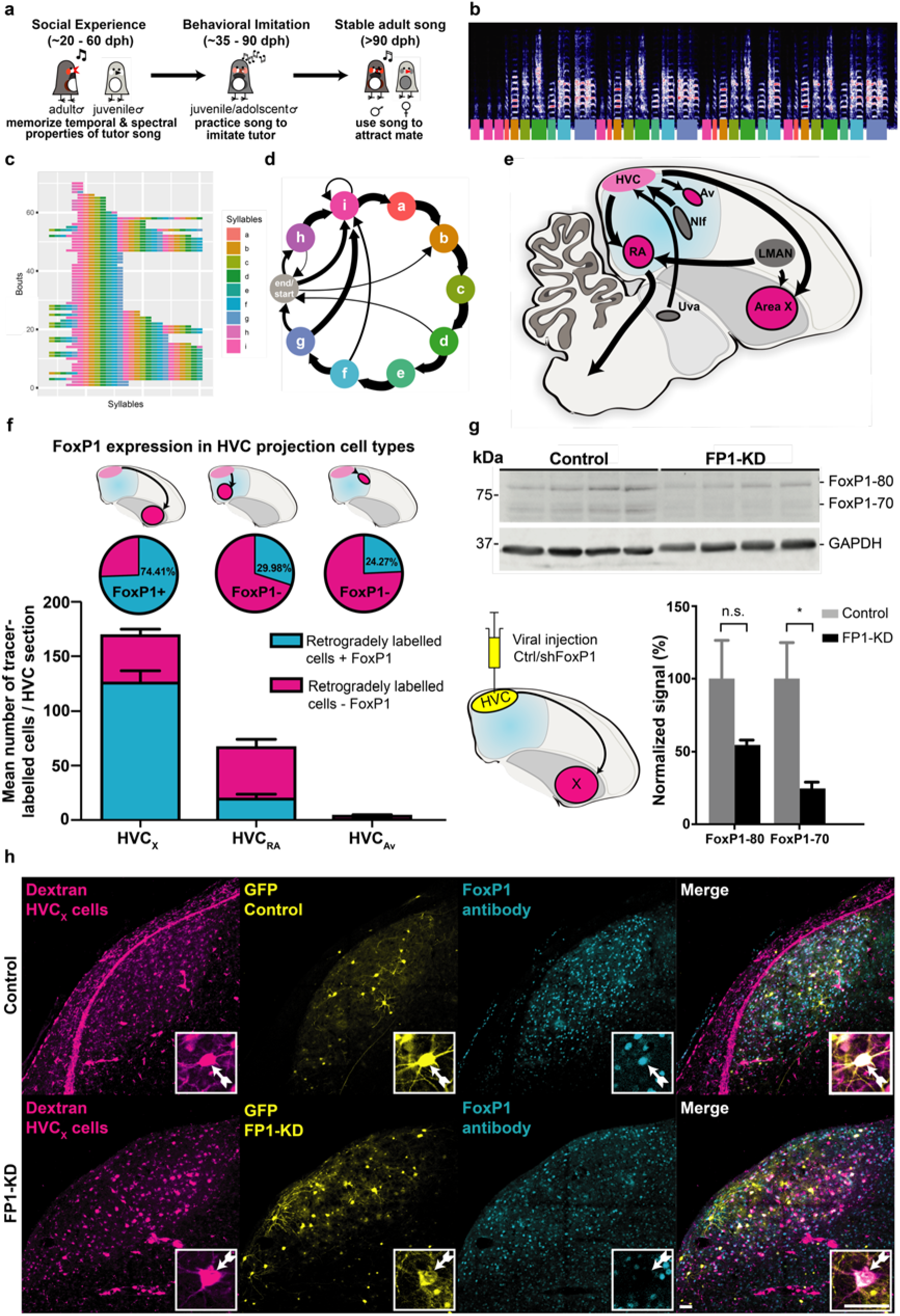
FoxP1 Expression in HVC of Control and Knock-down Zebra Finches. **a,** Timeline of zebra finch song learning in juvenile males (dph = days post hatch). **b-d,** Example representations of an adult zebra finch song; each color represents a syllable or note in the song: **b,** Spectrogram of an adult male’s song. Frequency is represented along the y-axis, time is represented along the x-axis, and color reflects intensity. Colored bars underneath indicate introductory notes (pink, i) and syllables (a-h). **c,** A syntax raster plot showing the syllables sung over repeated song bouts, colors reflect the syllables produced. **d,** A representation of song syllable syntax as a first-order Markov model, with thickness of arrows representing the probability of syllable transitions. **e,** Parasagittal schematic of the song circuit, with relevant nuclei labelled: Area X, striato-pallidal basal ganglia nucleus; Av, nucleus avalanche; HVC, premotor song nucleus; LMAN, lateral magnocellular nucleus of the anterior nidopallium; NIf, nucleus interfacialis of the nidopallium; Uva, nucleus uvaeformis; RA, robust nucleus of the arcopallium. **f,** FoxP1 expression in HVC projection neurons. Subtypes of HVC neurons, L-R: HVC_X_ neurons, HVC_RA_ neurons, HVCAv neurons. (top) Schematic of retrograde injections into areas downstream of HVC, (middle) pie charts showing the proportion of cells that express FoxP1 for each cell-type, and (bottom) proportion of FoxP1 expressing neurons for each HVC subtype, per HVC section (HVC_X_: 74.4±2.2%, *n*=3 birds, 6 hemispheres; HVC_RA_: 30.0±0.7%, *n*=2 birds, 4 hemispheres; HVCAv: 24.3±2.5%, *n*=3 birds, 6 hemispheres). **g,** Western blot of lysates from HVC injected with control or shFoxP1 AAV. At least two isoforms of FoxP1 were observed around 80 kDa and 70 kDa, with GAPDH as loading control. Schematic of injections for the tissues collected after viral injections of rAAV9/ds-CBh-GFP (*n*= 4 birds) or pscAAV-GFP-shFoxP1 (*n*= 4 birds) into HVC, paired with retrograde tracer in Area X. Graph shows quantification of FoxP1 protein. Signals were normalized to GAPDH, averaged for each condition, and normalized to the Controls. Histograms represent average±SEM. (<u>FoxP1-80</u>: Control: 100±26.6% vs FP1-KD: 54.7±3.5%, Student’s t-test, *P*> 0.05; <u>FoxP1-70</u>: Control: 100±25.0% vs FP1-KD: 24.7±4.4%, Student’s t-test, *P*= 0.025). **h,** Representative examples of HVC sections from Control (top) and FP1-KD (bottom) birds. Injections were performed as in schematic in panel ‘g’. Panels show HVC_X_ cells labelled with retrograde tracer in Area X (magenta, left), GFP signal from AAV-Control/ AAV-shFoxP1 injection (yellow, middle left), FoxP1 staining with antibody (cyan, middle right), and a merged composite (right). Inset boxes indicate one example cell per condition. White arrowheads indicate the soma of the example neurons. The FP1-KD cell (bottom insets) does not show FoxP1 labelling, whereas it is present in the Control cell (top insets). Scale bars, 50 μm.

FOXP1 (forkhead-box protein 1) is one of the top ASD-associated genes, and its haploinsufficiency causes specific language impairment and intellectual disability in children^5^. *FoxP1* is expressed in many of the same areas of the pallium and basal ganglia in mammals and songbirds^9–11^. In zebra finches, *FoxP1* expression is enriched in many brain regions that are known to be important for song learning^10–12^ (Fig. 1e, Extended Data Fig. 2). Here we focus on the role of FoxP1 in the pallial region HVC, which is involved in the formation of song memories, the vocal imitation process, and in production of the learned song^13–19^. We first characterized FoxP1 expression in HVC projection cell-types, then tested its necessity in different aspects of song learning. Lastly, we identified anatomical, electrophysiological and circuit abnormalities associated with disrupted expression of FoxP1 during vocal learning.

## FoxP1 is Expressed in Striatal Projecting HVC_X_ Neurons

HVC has three non-overlapping classes of projection neurons^20^ (Fig. 1e). HVC_X_ and HVC_Av_ neurons transmit vocal motor-related signals to the striatum or auditory system, respectively. These two pathways are important for behavioral imitation of tutor-song in juvenile birds, but are not essential for song production in adult birds^20–22^. HVC_RA_ neurons provide descending motor commands to the pallial nucleus RA, a connection which is necessary for the production of learned song at all stages of life^17,21,23^. We used anatomical tracing and immunolabeling to examine FoxP1 expression in these different classes of projection neurons. We found that FoxP1 is expressed in the majority of HVC neurons projecting to the striatum (74.41 ± 2.17% of HVC_X_ neurons) and in a much smaller percentage of neurons projecting to the auditory system or to RA (24.27 ± 2.54% of HVCAv neurons, 29.98 ± 0.65% of HVC_RA_ neurons, Fig. 1f, Extended Data Fig. 2a-f). The widespread expression of FoxP1 in striatal projecting HVC_X_ neurons raises speculation that FoxP1 could function in sensorimotor aspects of song learning. HVC_X_ neurons are thought to carry timing information about song to the basal ganglia, which facilitate accurate reinforcement-based imitation of syllables and sub-syllable elements^24–27^. In support of this view, genetic lesions of HVC_X_ neurons have recently been shown to significantly disrupt behavioral imitation of song in juvenile male zebra finches^22^.

## Knockdown of FoxP1 in HVC Blocks Learning from Social Experiences but not Behavioral Imitation of Song

To test the function of FoxP1 in the social transmission of birdsong we developed a short hairpin RNA (shRNA) against FoxP1 and showed that it can significantly knockdown expression of FoxP1 *in vivo* and *in vitro* (Fig. 1g-h, Extended Data Fig. 2h). Using an adeno associated virus to express this shFoxP1, we knocked-down expression of FoxP1 (FP1-KD) in age-matched juvenile birds prior to different aspects of song learning.

Juvenile male zebra finches memorize the song of their father or other adult song-tutor(s) during social interactions. They then use auditory feedback and extensive practice to learn how to accurately imitate this memorized song by 90100 days post hatching (dph)^18^. Juvenile birds can memorize the song of a tutor at any time between 20-60 dph, but they don’t start to practice singing until approximately 35-40 dph (Fig 1a). This developmental progression and the ability to raise birds in groups without a song-tutor – hereby referred to as ‘isolates’– allowed us to knock down *FoxP1*expression prior to behavioral imitation of song, either before or after birds had an opportunity to form a memory of a tutor-song^16^ (Fig. 2a: tutoring pre-FP1-KD, referred to as ‘behavioral imitation’ group (**FP1-KD BI**), Fig.2e: tutoring post-FP1-KD, referred to as ‘social experience’ group (**FP1-KD SE**)). Given the widespread expression of FoxP1 in striatal projecting HVC_X_ neurons, we hypothesized that FP1-KD might disrupt behavioral imitation of the tutors’ song^22^. However, we found that FP1-KD after birds had an opportunity to form a memory of the tutor-song (FP1-KD BI birds) did not disrupt their ability to then copy that song during development (Fig. 2a-d). In fact, the adult songs of the FP1-KD BI group were stereotyped and indistinguishable from normal zebra finch song, suggesting that FoxP1 in HVC is not necessary for behavioral imitation in juvenile birds or in more basic aspects of song production.

**Figure 2.**
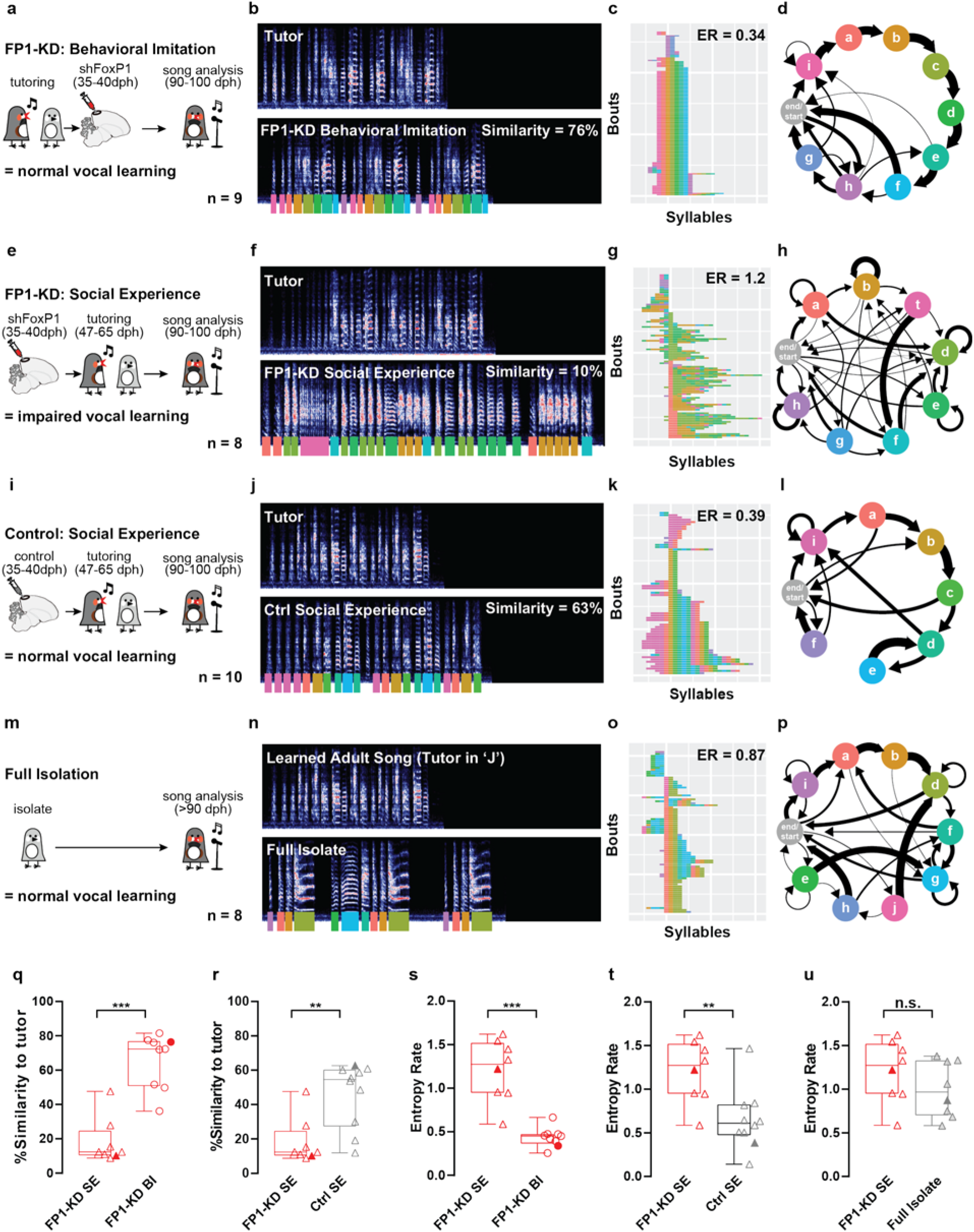
Song learning is impaired by FoxP1 KD. **a, e, i** & **m,** Timelines illustrating tutoring experience of FP1-KD behavioral imitation (**a**), FP1-KD social experience (**e**), Control social experience (**i**), and full isolate birds (**m**). **b, f, j** & **n**, Representative spectrograms from a single bird belonging to each experimental group and their tutor. Colored underlines reflect syllable labels used for subsequent syntax visualizations and analysis. Similarity refers to % similarity to tutor, also represented in **q** & **r**. The full isolate bird (**n**) had no tutor, and therefore has no similarity to tutor score. It is contrasted with the song of an adult zebra finch with typical tutor exposure. **c, g, k** & **o**, Syntax raster plot illustrating the syntax stereotypy of an example bird for each condition. This is the same bird used for the spectrogram and syntax diagram. Each row reflects a single song bout, and each colored block reflects the syllable sung at that position in the bout. Rows are sorted according to syllable order. ER refers to entropy rate, also represented in **s-u. d, h, l** & **p,** Diagrams reflecting syllable transitions produced by an example bird for each condition. Line thickness is proportional to the transition probability from the originating syllable to the following. Transitions with a probability of less than 4% are omitted for clarity. **q.** FP1-KD social experience birds (*n*=8) have significantly lower song similarity to tutor than FP1-KD behavioral imitation birds (*n*=9, FP1-KD SE: 12.39 vs FP1-KD BE: 72.32, Mann-Whitney test, *P* < 0.001). Filled points correspond to the example birds shown in **f-h** and **b-d. r**, FP1-KD social experience birds (*n*=8) have significantly lower song similarity to tutor than Control social experience birds (*n*=10, FP1-KD SE: 12.39 vs Ctrl SE: 54.6, Mann-Whitney test, *P* = 0.0031). Filled points correspond to the example birds shown in **f-h** and **j-l. s**, FP1-KD social experience birds (*n*=8) have significantly higher song syntax entropy rates than FP1-KD behavioral imitation birds (*n*=9, FP1-KD SE: 1.274 vs FP1-BI: 0.4512, Mann-Whitney test by bird, *P* < 0.001). Filled points correspond to the example birds shown in **f-h** and **b-d. t**, FP1-KD social experience birds (*n*=8) have significantly higher song syntax entropy rates than Control social experience birds (*n*=10, FP1-KD SE: 1.274 vs Ctrl SE: 0.6113, Mann-Whitney test, *P* = 0.0085). Filled points correspond to the example birds shown in **f-h** and **j-l. u**, Song syntax entropy rates do not differ significantly between FP1-KD social imitation (*n*=8) and full isolate birds (*n*=8, FP1-KD SE: 1.274 vs Full Isolate: 0.9667, Mann-Whitney test, *P* > 0.05). Filled points correspond to example birds shown in **f-h** and **n-p**. For all box plots, median, 25^th^ and 75^th^ percentile, and minimum and maximum are reported. The single datapoints are overlaid on the side.

In contrast, FP1-KD prior to social experience with a tutor severely disrupted subsequent song learning (FP1-KD SE birds, Fig. 2e-h). We found that birds with FP1-KD prior to social experience learned little from their song-tutor and statistically less than birds in the FP1-KD BI or control birds (Fig. 2a-l, q-r). As adults, FP1-KD SE birds sang songs that were highly variable from trial-to-trial and with entropy rates (Methods) higher than either the FP1-KD BI or control birds (Fig. 2a-l, s-t). Rather, their songs had entropy rates that were statistically indistinguishable from birds that were never tutored during development (full isolates, Fig. 2 m-p, u). Together, these findings suggest that FP1-KD critically impairs the cultural transmission of vocal behavior, potentially by selectively disrupting the ability to form appropriate vocal memories during social experiences.

## Knockdown of FoxP1 Inhibits Dendritic Spine Turnover and Decreases the Intrinsic Excitability of HVC_X_ Neurons

We next sought to identify the consequences of FP1-KD on structural and functional plasticity in HVC. Previous research has shown that plasticity in HVC is predictive of a young bird’s ability to form tutor-song memories during social interactions^13,14^. Birds with high levels of dendritic spine turnover are better learners than birds with low levels of spine turnover^14^. Therefore, we first used longitudinal *in vivo* two-photon imaging to track FP1-KD mediated changes to dendritic spines in our social experience groups and in separate cohorts of juvenile isolates (Fig. 3a). We imaged spines on virally transduced, retrogradely labeled HVC_X_ neurons, both in our FP1-KD cohorts and in control birds. We found that HVC_X_ neurons had significantly higher spine density in FP1-KD adults and lower spine density in FP1-KD juveniles than in their respective age-matched controls (Fig. 3b-c). This suggests that FP1-KD disrupts normal changes in spine density that occur during song learning. Perhaps more pertinently, we found that FP1-KD signifcantly reduced spine turnover in both juvenile and adult birds compared to age-matched control birds (Fig. 3d-e). We then verified these results with independent analysis by 4 additional scorers on a randomized subset of this data (*P* < 0.05, Paired t-test). Therefore, plasticity in the striatal projecting HVC_X_ neuronal population may be necessary for young birds to learn from social experiences and to form the vocal memories used to guide song imitation.

**Figure 3.**
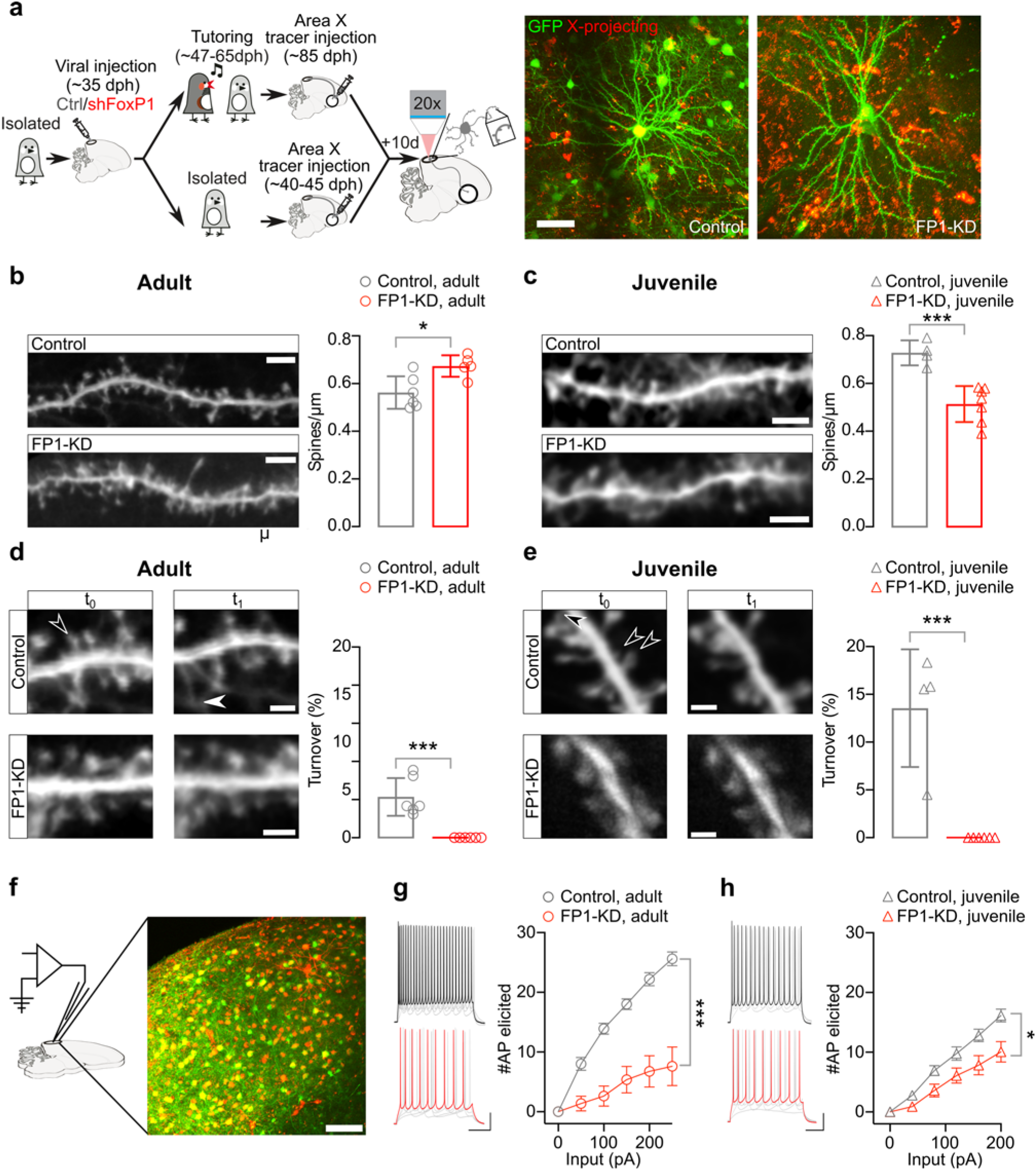
FP1-KD reduces structural plasticity and excitability of HVC_X_ neurons. **a,** (left) schematic of the experimental protocol and timeline of the experiments. (right) *In-vivo* two photon images of sample GFP-labelled (green) and retrogradely labelled (red) Control and FP1-KD HVC_X_ neurons. Scale bar, 50 μm. **b,** (left) Representative *in-vivo* two photon images of GFP-expressing dendrite sections from Control (top) and FP1-KD (bottom) normally reared adult HVC_X_ neurons. Scale bar, 5 μm. (right) Histograms reporting the average±SEM dendritic spine density (spines/μm) of Control and FP1-KD dendrites from adult HVC_X_ neurons (Control adult: 0.56±0.03, *n*=821 spines, 6 cells, 2 animals; FP1-KD adult: 0.67±0.02, *n*=668 spines, 6 cells, 5 animals; Student’s t-test, *P* = 0.01). **c,** (left) Representative *in-vivo* two photon images of GFP-expressing dendrites sections from Control (top) and FP1-KD (bottom) juvenile isolate HVC_X_ neurons. Scale bar, 5 μm. (right) Histograms reporting the average±SEM dendritic spine density (spines/μm) of Control and FP1-KD dendrites from juvenile HVC_X_ neurons (Control juvenile: 0.73±0.03, *n* =769 spines, 4 cells, 3 animals; FP1-KD juvenile: 0.51±0.03, *n*=745 spines, 6 cells, 5 animals; Student’s t-test, *P* < 0.001). **d,** (left) Representative images of Control and FP1-KD adult dendritic segments from HVC_X_ neurons taken at two different times (t_0_,t_1_ across a 4-h imaging interval). Filled arrowhead indicates a representative gained synapse. Empty arrowhead indicates a representative lost synapse. Scale bars, 2 μm. (right) Histograms representing the average±SEM percent dendritic spine turnover (acquired + lost spines/total spines counted) from Control and FP1-KD adults (Control adult: 4.3±0.8%, *n*=1126 spines, 6 cells, 2 animals; FP1-KD adult: 0.0±0.0%, *n*=1148 spines, 6 cells, 5 animals; Student’s t-test, *P* < 0.001). **e,** (left) Representative images of Control and FP1-KD juvenile dendritic segments from HVC_X_ neurons, taken at two different times (t0,t1 across a 2-h imaging interval). Empty arrowhead indicates a representative lost synapse. Scale bars, 2 μm. (right) Histograms representing the average±SEM percent dendritic spine turnover (acquired + lost spines/total spines counted) from Control and FP1-KD juveniles (Control juvenile: 13.6±3.1%, *n*=650 spines, 4 cells, 3 animals; FP1-KD juvenile: 0.0±0.0%, *n*=735 spines, 6 cells, 5 animals; Student’s t-test, *P* < 0.001). **f,** Schematic of an *ex-vivo* slice and patch-clamp recording setup with high-resolution image of HVC in a brain slice used for electrophysiology. Scale bar, 50 μm. g, Example traces (left, scalebars 20 mV, 100ms) and plot (right) reporting the number of action potentials elicited by somatic current injections in HVC_X_ neurons from Control and FP1-KD adult brain slices. FoxP1 knockdown decreased the intrinsic excitability of HVC_X_ neurons in adults (Controls, *n* = 10, 5 animals; FP1-KD *n*= 8, 3 animals; Two-Way Anova, Interaction F_10,160_ = 30.87, Treatment F_1,16_ = 34.56, *P* < 0.001). h, Example traces (left, scalebars 20 mV, 100ms) and plot (right) reporting the number of action potentials elicited by somatic current injections in HVC_X_ neurons from Control and FP1-KD juvenile brain slices. FoxP1 knockdown decreased the intrinsic excitability of HVC_X_ neurons in isolate juveniles (Controls, *n*=12, 3 animals; FP1-KD *n*=6, 2 animals; Two-Way Anova, Interaction F_10,200_ = 7.053, Treatment F_1,20_ = 7.627, *P* = 0.01). All data are reported as average±SEM. In the histograms, the single datapoints are overlaid on the side of each bar.

This decreased spine turnover in FP1-KD birds could be tied to differences in the intrinsic excitability of HVC_X_ neurons. For example, deafening of adult zebra finches causes increased intrinsic excitability and increased dendritic spine turnover on HVC_X_ neurons^28^. To examine this, we conducted whole-cell current-clamp recordings of HVC_X_ neurons from our social experience groups (FP1-KD and control) and from cohorts of juvenile isolates (Fig. 3f). We found that neurons expressing shFoxP1 had significantly decreased intrinsic excitability compared to HVC_X_ neurons recorded from age-matched controls (Fig. 3g-h).

## Knockdown of FoxP1 Blocks Synaptic and Network Hallmarks of Learning during Social Experience

We next explored if social experience with a tutor, and presumably formation of a vocal memory, is sufficient to elicit changes in the intrinsic excitability and synaptic physiology of HVC_X_ neurons. We made comparisons between juvenile isolates and age-matched birds that were housed with a song tutor for two consecutive days (Fig. 4a). We found that this acute social experience with a tutor drove a significant increase in the intrinsic excitability of HVC_X_ neurons in control birds. FP1-KD prevented these changes in HVC_X_ neurons (Fig. 4b). With regards to synaptic properties, we found that neither FP1-KD nor social experience affected the excitation-to-inhibition ratio (E/I ratio) in HVC_X_ neurons (Fig. 4c). However, social experience led to a significant increase in AMPA/NMDA receptor ratios in HVC_X_ neurons from control birds (Fig. 4d), a result consistent with synaptic strengthening following tutor experience and the known dependence on NMDA receptor activity in HVC for encoding of tutor song memories^13^. In contrast, FP1-KD prevented this AMPA/NMDA increase following social experience. These results indicate that FP1-KD blocks key synaptic and intrinsic electrophysiological signatures of learning following social experiences with a song-tutor.

**Figure 4.**
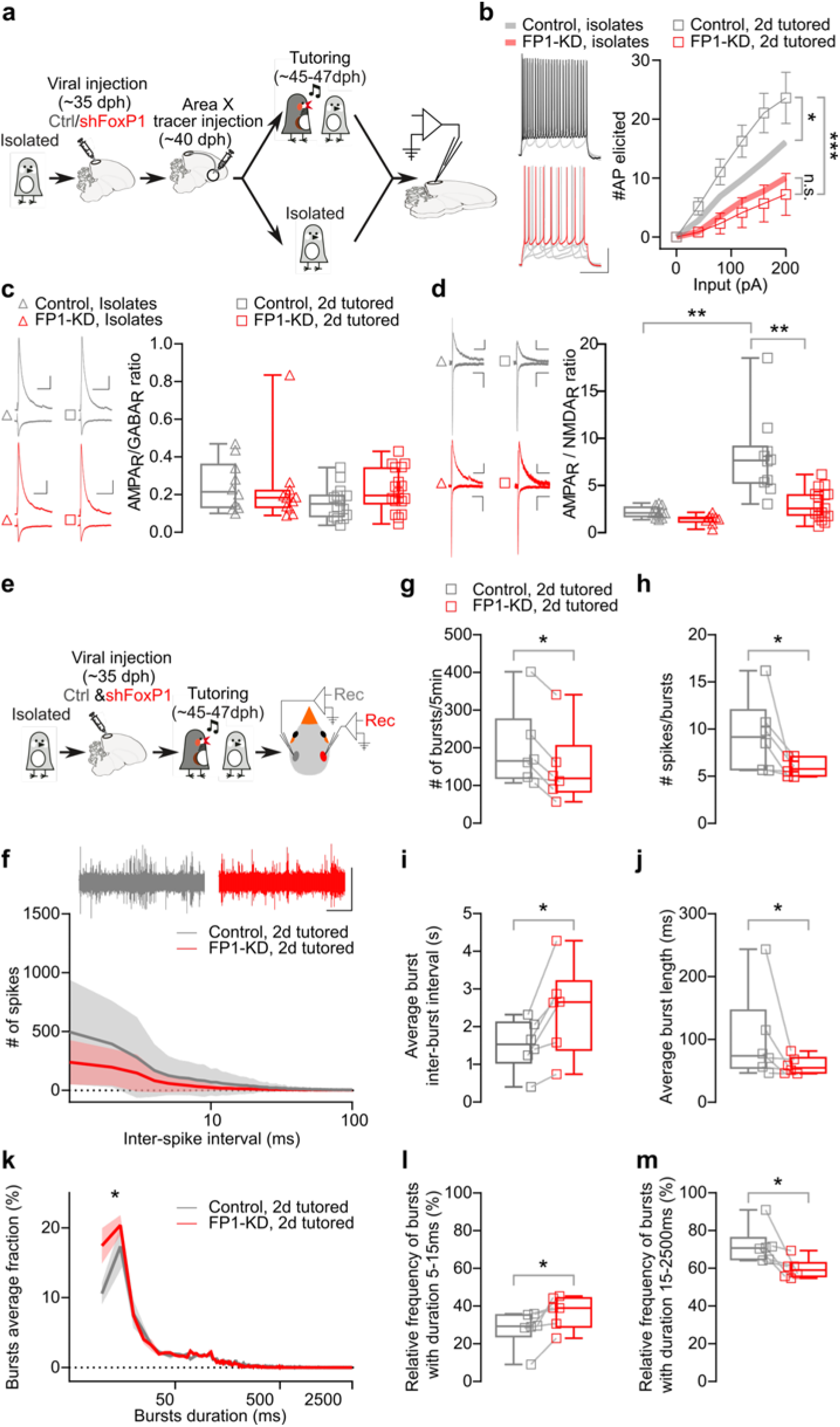
FP1-KD prevents experience-dependent synaptic strength modifications and reduces network-level bursting activity organization. **a,** Experimental timeline. **b,** Example traces (left) and plot (right) describing the number of spikes elicited by somatic depolarization steps in 2-day tutored Control and FP1-KD HVC_X_ neurons. Trendlines relative to the same experiments, but conducted in isolates, are reported here from Fig 3h. HVC_X_ neurons were more excitable in control birds subjected to the 2-day tutoring regime compared to isolates (Two-Way Anova, Interaction F_10,190_ = 4.598, Treatment F_1,19_ = 5.376, *P* = 0.03). This difference is not present in FP1-KD birds (Two-Way Anova, Interaction F_10,180_ = 0.4578, *P* > 0.05). Among 2 day-tutored birds, FoxP1 knockdown resulted in a decreased intrinsic excitability of HVC_X_ neurons (Controls, *n*= 9, 4 animals; FP1-KD *n*= 10, 3 animals; Two-Way Anova, Interaction F_10,170_ = 9.308, Treatment F_1,17_ = 10.73, *P* = 0.005). Data are reported as average±SEM. **c,** Representative traces (left), box and scatter plots (right) of evoked AMPAR and GABAR mediated currents recorded at −60 mV or 10 mV respectively (scale bars:100 pA; 100 ms) in Control or FP1-KD HVC_X_ neurons from isolate and 2-day tutored birds. Box plots represent the AMPAR/GABAR currents amplitude ratios from isolates (open triangles) and 2-day tutored birds (open squares). FoxP1 knockdown has no significant effect on the AMPAR/GABAR ratios for both bird groups (Isolates Control: 0.25±0.04, *n*=10, 5 animals; isolated FP1-KD: 0.23±0.06, *n*=11, 5 animals; 2d-tutored Control: 0.16±0.02, *n*=14, 5 animals; 2d-tutored FP1-KD: 0.22±0.03, *n*=18, 6 animals; Kruskal-Wallis test, *P* > 0.05). **d,** Representative traces (left), box and scatter plot (right) of evoked AMPAR and NMDAR mediated currents recorded at −70 mV or +40 mV respectively (scale bars:50 pA; 100 ms) in Control or FP1-KD HVC_X_ neurons in isolates and 2-day tutored birds. Box plots represent the AMPAR/ NMDAR current ratios recorded from isolates (open triangles) and 2-day tutored birds (open squares). (isolate Control: 2.2±0.2, *n*=12, 6 animals; isolate FP1-KD: 1.4±0.2, *n*=8, 5 animals; 2d-tutored Control: 8.0±1.4, *n*=10, 6 animals; 2d-tutored FP1-KD: 2.9±0.4, *n*=17, 6 animals; Kruskal-Wallis test, *P* < 0.001; Dunn’s multiple comparisons, isolate Control vs 2d-tutored Controls, *P* = 0.002, isolate FP1-KD vs 2d-tutored Controls, *P* < 0.001, 2d-tutored Controls vs 2d-tutored FP1-KD, *P* = 0.008). **e,** Schematic representing the experimental timeline. Birds received AAV injections in HVC to knock-down FoxP1 in one hemisphere, and control virus in the other hemisphere (pseudorandomized). The birds were kept in isolation from male tutors, and ~10 days later they received 48h tutoring, after which HVC extracellular activity was recorded in both hemispheres (*n* = 6 birds, 3-5 recordings/hemisphere). **f,** (top) Sample traces (scalebar 0.5V, 1s) and (bottom) average inter-spike interval distribution (bin 1ms, 1-100ms, logarithmic scale, 300s/recording) (Control hemispheres vs FP1-KD hemispheres, Two-Way Anova, Interaction F99,990 = 1.222, *P* > 0.05). Data are reported as average (thick line)±SEM (semitransparent contour) **g,** Box and scatter plot of the total number of bursts (Control hemispheres: 199.4 ± 44.4 vs FP1-KD hemispheres: 148.0 ± 41.2, Wilcoxon matched-pairs signed rank test, *P* = 0.03). **h,** Box and scatter plot of the average number of spikes in a burst (Control hemispheres: 9.4 ± 1.6 vs FP1-KD hemispheres: 6.0 ± 0.4, Wilcoxon matched-pairs signed rank test, *P* = 0.03). **i,** Box and scatter plot of the average inter-burst interval (Control hemispheres: 1.5 ± 0.3 vs FP1-KD hemispheres: 2.4 ± 0.5, Wilcoxon matched-pairs signed rank test, *P* = 0.03). **j,** Box and scatter plot of the average burst length (Control hemispheres: 101.6 ± 30 vs FP1-KD hemispheres: 58.6 ± 5.8, Wilcoxon matched-pairs signed rank test, *P* = 0.03). **k,** Plot representing the relative distribution of burst duration, normalized for each recording (5ms duration bins, 5-2500ms, logarithmic scale; Control hemispheres vs FP1-KD hemispheres, Two-Way Anova, interaction F_498,4980_ = 1.972, *P* < 0.001, Control vs FP1-KD F_1,10_ = 5.570, *P* = 0.04). Data are reported as average (thick line)±SEM (semitransparent contour). **l,** Box and scatter plot of the average relative prevalence of bursts with durations between 5 and 15ms (Control hemispheres: 27.9 ± 4.0 vs FP1-KD hemispheres: 36.8 ± 3.5, Wilcoxon matched-pairs signed rank test, *P* = 0.03). **m,** Box and scatter plot of the average relative prevalence of bursts with durations between 15 and 2500ms (Control hemispheres: 72.1 ± 4.0 vs FP1-KD hemispheres: 59.8 ± 2.2, Wilcoxon matched-pairs signed rank test, *P* = 0.03). For all box plots, median, 25^th^ and 75^th^ percentile, and minimum and maximum are reported. The single datapoints are overlaid on the side.

Lastly, we tested whether these cellular and synaptic effects of FP1-KD were sufficient to block *in vivo*, network-level hallmarks of song memory. Acquisition of tutor song memories following brief social experiences is correlated with the emergence of prolonged patterns of bursting activity in HVC and in RA^14,15,29^. Taking advantage of the lack of a corpus callosum connecting HVC from the right and left hemispheres^30^, we conducted experiments in juvenile isolates in which we knocked down FoxP1 in only one, pseudo-randomly assigned, brain hemisphere and expressed a control virus in the other. Birds were either maintained in social isolation from a song-tutor (Extended Data Fig. 4a) or given two days of social experience with a song-tutor (Fig. 4e), before making bilateral extracellular recordings from HVC to assess baseline and learning related changes in network activity. In isolate birds (baseline condition), we did not detect any network level differences in spontaneous neuronal activity between the FP1-KD and control hemispheres (Extended Data Fig. 4 and 5a-b). This indicates that FP1-KD does not cause large scale changes in the excitability or bursting properties of HVC neurons in the absence of prior tutor-song experience.

In contrast, following two-days of social experience with a tutor, we observed large scale differences in the spontaneous bursting activity recorded in the control and FP1-KD hemispheres. We observed significantly fewer bursts in the FP1-KD hemispheres, and these bursts had fewer spikes and were shorter than bursts recorded in the control brain hemispheres (Fig. 4g-m). The total number of spikes, inter-spike interval distribution, and inter-spike interval within bursts were not affected by FP1-KD (Fig. 4f and Extended Data Fig. 5c-d). This suggests that while overall activity levels were preserved in the two hemispheres, the tutor-song experience drove redistribution of activity into sustained bursting patterns in the control, but not in the FP1-KD hemisphere. This indicates that knockdown of FoxP1 is sufficient to block circuit level hallmarks of learning from social experiences and encoding of vocal memories that guide the cultural transmission of song.

## Discussion

How ASD risk genes differentially impact learning of culturally transmitted behaviors is difficult to study in traditional model organisms; many don’t robustly imitate social behaviors nor transmit behavioral repertoires from one generation to the next^31^. Our results demonstrate that FP1-KD produces a series of synaptic, cellular and network deficits, some of which resemble those recently described in mouse models^32–34^. However, the characteristics of the songbird system allowed us to test the function of FoxP1 in social experience-dependent imitation of behavior. We show that FoxP1 in HVC is essential for young birds to encode memories that are used to guide imitation, but not for the imitation of a previously acquired song memory. Identified roles of HVC include motor control of song, vocal imitation, and encoding of tutor-song memories^13–20^. Therefore, FP1-KD mediated selective disruption of learning from social experiences is surprising and indicates a specialized and essential role for the network of FoxP1-expressing HVC neurons in the cultural transmission of vocal behaviors.

Notably, we find that FoxP1 is widely expressed in striatal projecting HVC_X_ neurons. These neurons have previously been shown to have properties resembling ‘mirror’ neurons^35–38^, but their role in the cultural transmission of vocal behaviors was not well understood. The functional significance of mirror neurons in behavioral imitation and ASD is a topic of much debate^39–44^. We show that knockdown of FoxP1 in striatal projecting HVC_X_ neurons inhibits their spine plasticity, dampens their intrinsic excitability, and blocks the cellular and network level signatures associated with tutor-song memory formation following social experience. Although it is not yet clear whether the disruptions in learning shown here depend exclusively on HVC_X_ neurons, as they are not the only cell-type in HVC expressing FoxP1, our research provides proof that a detailed examination of how ASD-linked genes affect different aspects of vocal imitation can offer novel and important insights into their roles in neurodevelopmental disorders.

## Acknowledgments

The authors thank members of the Roberts and Konopka laboratories for discussion and comments on the manuscript, Andrea Guerrero and Matthew Harper for laboratory support.

## Funding

This research was supported by grants from the US National Institutes of Health R21DC016340 to TFR and GK, R01NS108424 & R01DC014364 to TFR, R01MH102603 to GK, and from the National Science Foundation IOS-1457206 to TFR. FA and DHA were supported by T32HL139438.

## Author contributions

TFR and GK conceived the project. TFR supervised the research. FG-O, TK, HP, MT and TFR designed the experiments and wrote the manuscript with input from all authors. FG-O, TK, HP, MT, VD, MC, DHA and JEH collected and analyzed the data for the project. GK, MC and FA designed and tested the knockdown viral construct. All authors read and commented on the manuscript.

## Competing interests

Authors declare no competing interests.

## Data and materials availability

All data is available in the main text or the supplementary materials.

## Supplemental Information

Materials and Methods

Extended Data Figures 1 to 5

## Materials and Methods

### Animals

Experiments described in this study were conducted using juvenile and adult male zebra finches (30-110 days post hatch (dph)). We raised juvenile male zebra finches either in isolation from an adult song model (isolates), with normal access to an adult song model (non-isolates), or isolates that were exposed to 2 days of tutoring by an adult song model (2-day tutored). All procedures were performed in accordance with protocols approved by Animal Care and Use Committee at UT Southwestern Medical Center.

### Viral Vectors

The following adeno-associated viral vectors were used in these experiments: rAAV2/9/ds-CBh-GFP (The University of North Carolina at Chapel Hill Gene Therapy Center Vector Core), and pscAAV-GFP-shFoxp1 (IDDRC Neuroconnectivity Core at Baylor College of Medicine). All viral vectors were aliquoted and stored at −80°C until use.

### Constructs

pscAAV-GFP-shFoxP1 was generated by PCR-amplification of U6-shFoxp1 from pLKO.1 (TRCN0000072005, Broad Institute) while adding NotI and BamHI sites, then ligating into pscAAV9-CBh-GFP (Xiao et al., 2018) digested with these enzymes. The PCR program used was: 98°C 2 min, 35 x (98°C 10 s, 55°C 15 s, 72°C 5 s), 72°C 7 min. The primers used were forward (5’-3’): ATAAGAATGCGGCCGCTTTCCCATGATTCCTTC, and reverse (3’-5’): CGCGGATCCAAAAAGCTAACACT AAACG. The scrambled control (shCtrl) was designed by uploading shFoxp1 into https://www.genscript.com/ssl-bin/app/scramble and selecting the chicken mRNA pool. Ensembl BLAST found only 3 hits of 15/21 bases in the zebra finch genome. pCMV-FoxP1^WT^ was generated by subcloning zebra-finch specific FoxP1 cDNA from pGEMTeasy (Haesler et al., 2004) into pCMV-Tag4a using EcoRI.

### Stereotaxic Surgery

All surgical procedures were performed under aseptic conditions. Birds were anaesthetized using isoflurane inhalation (0.8-1.5%) and placed in a stereotaxic surgical apparatus. The centers of HVC and RA were identified with electrophysiological recordings, and Area X and Av were identified using stereotaxic coordinates.

Viral injections to HVC were performed using previously described procedures (Roberts et al., 2012; Roberts et al., 2017). Briefly, adeno-associated viral vectors (pscAAV-GFP-shFoxP1 or rAAV9/ds-CBh-GFP) were injected into HVC (50 nl per injection and ~60 injections, for a total of ~3 μl) at ~35 dph, and the transgenes were allowed to express for a minimum of 10 days. We also injected 500-950nl of differently conjugated tracers (Dextran, AlexaFluor 488 or 594, 10,000MW, Invitrogen) bilaterally into birds’ Area X, Av, and RA respectively. Tracer injections were performed using previously described procedures (Roberts et al., 2012; Roberts et al., 2017; Xiao et al., 2018) at the following approximate stereotaxic coordinates relative to interaural zero and the brain surface (rostral-caudal, medial-lateral, dorsal-ventral, in mm, head angle): HVC (0, ±2.4, 0.1-0.6, with 30° head angle); RA (−1.0, ±2.4, 1.7-2.4, with 30° head angle); Area X (5.1, ±1.6, 3.3, with 45° head angle), and Av (1.75, ±2.0, 1.0, with 45° head angle).

### Tutoring Conditions

<u>Social Experience (SE)</u>: juvenile male zebra finches raised in isolation from an adult song model were injected with viruses into HVC at 35-40 dph. After optimal viral expression, these juveniles were then housed with a song tutor between days ~47 and 65 dph. All birds were separated from their tutors at 65 dph and raised to adulthood.

<u>Behavioral Imitation</u> (BI): juvenile male zebra finches were reared with a song tutor and injected with viruses into HVC at 35-40 dph. Normally reared juveniles were raised with an adult song model before and after viral injections, separated from their tutors at 60 days of age, and raised to adulthood.

<u>2-day Tutored</u>: juvenile male zebra finches that were raised as isolates were injected with viruses into HVC at ~35 dph. After allowing time for viral expression, a song tutor was placed into the isolate’s cage for two days of tutoring. Birds were then separated from their tutors and raised to adulthood.

<u>Isolates</u>: juvenile male zebra finches raised in isolation from an adult song model were injected with viruses into HVC at ~ 35 dph, and raised to adulthood without exposure to an adult song model.

### *In vivo* two-photon imaging

We conducted longitudinal dendritic spine imaging in male juvenile zebra finches raised in isolation from an adult song model (isolates) aged 45–55d or adult zebra finches 90-100d (table details). Viruses (pscAAV-GFP-shFoxP1 or rAAV9/ds-CBh-GFP) were allowed to express for a minimum of 14 days before a cranial window over HVC was made. Birds were anaesthetized by isoflurane inhalation (2%) and positioned in a stereotaxic apparatus for cranial windowing. The scalp overlying HVC was removed and the scalp margins were sealed to the surface of the skull using Vetbond (*n*-butyl cyanoacrylate). Bilateral craniotomies (~1-1.5 mm^2^) were made in the skull overlying HVC. The dura mater was excised, leaving the pia mater, the 60–150-μm-thick layer of neural tissue, and the lateral telencephalic ventricle overlying HVC intact. A custom-cut coverslip (no.-1 thickness) was placed directly on the pial surface, then sealed to the skull with dental acrylic. A head post was also affixed to the skull with dental acrylic as per previously described procedures (Daliparthi et al., 2019). Birds were placed onto a custom stage under an Ultima IV Bruker laser scanning microscope running Prairie View Software. Only HVC neurons that expressed both the retrograde tracer Dextran Alexa Fluor 594 from Area X (HVC_X_ neurons) and GFP from viruses were chosen for spine imaging. Dendritic segments of these neurons were imaged at high resolution during the bird’s subjective night-time (1024 × 1024 pixels, 76 × 76 μm^2^, 3.2 μs per pixel, averaging two samples per pixel with 1-μm *z* steps, focused through a ×20, NA 1.0 Zeiss IR-Achroplan immersion objective). Birds were then returned to a darkened holding cage and allowed to sleep, and were re-imaged 2-4 hrs later.

### Spine Image Analysis

Dendritic spine images were analyzed as reported previously (Roberts et al., 2010).Briefly, three-dimensional image stacks were auto-aligned and smoothed using a Gaussian filter (ImageJ; http://rsbweb.nih.gov/ij/) and the same dendritic segment, imaged twice with a 2 hr interval, was selected. Images exhibiting changes in fluorescence or rotational artefacts were excluded from further analysis. Sets of selected three-dimensional image stacks were scored by 4 researchers blind to the experimental condition. To assess spine growth and retraction, we compared individual dendritic spines across 2-4 hr time intervals and calculated spine stability (*N*_stable_/*N*_total_), spine elimination (*N*_lost_/*N*_total_), spine addition (*N*_gained_/*N*_total_) and spine turnover ((*N*_gained_ + *N*_lost_)/2*N*_total_), where *N*_stable_ is the number of spines that were stable over the time interval, *N*_lost_ is the number of spines lost over the time interval, *N*_gained_ is the number of spines gained over the time interval and *N*_total_ is the total number of spines from the first imaging time point. Changes in spine density (*N*_total_ divided by dendritic length in μm) were measured from the same dendritic segments used to assess spine turnover.

### *In-vivo* extracellular recordings

Isolated, juvenile (dph 30-35) birds received a unilateral (pseudo-randomized) injection of pscAAV-GFP-shFoxp1 in HVC. The other hemisphere received the control virus (rAAV9/ds-CBh-GFP). After 10-12 days, we recorded HVC electrophysiological activity from both hemispheres (3-5 recordings/hemisphere, minimum 5 minutes/recording after an initial 5 minutes to allow the activity to stabilize after the electrode dip). Recordings from both HVCs were sampled in a pseudorandomized order. Signals were acquired at 10KHz, and filtered (high-pass 300Hz, low-pass 20KHz). We used Spike2 to analyze the spikes whose amplitude reached a threshold of 0.3V (determined based on the average noise level among all the recordings). A Spike2 script was used to analyze the characteristics of spikes in bursting patterns. Bursts were defined as a minimum of 2 spikes separated by 10ms or less, and the burst epoch was considered terminated if no spike was detected for 100ms after the last spike of the burst. Values for each recording were averaged per hemisphere, and statistically compared pairwise where indicated to reveal differences. For the comparison of bursts length, the data from each hemisphere was normalized to produce a frequency distribution of the burst lengths for that hemisphere (bin size 1ms). The data were then compared across treatments, and further subdivided into two burst duration categories: 5-15ms, and 15-2500ms.

### *Ex vivo* physiology

#### Slice preparation

All extracellular solutions were adjusted to 310 mOsm, pH 7.3-7.4, and aerated with a 95% O_2_/5% CO_2_ mix. Zebra finches were first deeply anesthetized with isoflurane. Once the bird was no longer responsive to a toe pinch, it was quickly decapitated. The brain was removed from the skull and submerged in cold (1– 4°C) oxygenated dissection buffer. Acute sagittal 230-250 μm brain slices were cut in dissection buffer at 4°C containing (in mM): 225 sucrose, 3 KCl, 1.25 NaH_2_PO_4_, 26 NaHCO_3_, 10 D-(+)-glucose, 2 MgSO4, .5 CaCl_2_, 2 Kynurenic acid. Individual slices were incubated in a custom-made holding chamber saturated with 95% O_2_/5% CO_2_ at 34 °C for 20 minutes, and then kept at 30 °C for a minimum of 45 minutes in artificial cerebrospinal fluid (aCSF), containing (in mM): 126 NaCl, 3 KCl, 1.25 NaH_2_PO_4_, 26 NaHCO_3_, 10 D-(+)-glucose, 2 MgSO_4_, 2 CaCl_2_.

#### Slice electrophysiological recording

Slices were constantly perfused in a submersion chamber with 32°C oxygenated normal aCSF. Patch pipettes were pulled to a final resistance of 3-5 MΩ from filamented borosilicate glass on a Sutter P-1000 horizontal puller. HVC_X_ cell bodies double-labelled with GFP and Dextran AlexaFluor594 were visualized by epifluorescence imaging using a water immersion objective (×40, 0.8 numerical aperture) on an upright Olympus BX51 WI microscope, with video-assisted infrared CCD camera (Q-Imaging Rolera). Data were low-pass filtered at 10 kHz and acquired at 2 kHz with an Axon MultiClamp 700B amplifier and an Axon Digidata 1550B Data Acquisition system under the control of Clampex 10.6 (Molecular Devices).

For **voltage clamp** whole-cell recordings of HVC_X_ projecting neurons, the internal solution contained (in mM): 120 Cs-gluconate, 10 HEPES, 5 tetraethylammonium-Cl, 2.8 NaCl, 0.6 EGTA, 4 MgATP, 0.3 NaGTP, 5 BAPPTA, 7 QX314 chloride (adjusted to pH 7.3-7.4 with CsOH, 297 mOsm). For **current clamp** recordings, the internal solution contained: 116 K-gluconate, 20 HEPES, 6 KCl, 2 NaCl, 0.5 EGTA, 4 MgATP, 0.3 NaGTP, 10 Na-phosphocreatine (adjusted to pH 7.3-7.4 with KOH, 299 mOsm).

Electrically evoked **synaptic** currents were measured by delivering one electric stimulus (1 ms, 10–30 μA) every 12 s, with an isolation unit, through a glass stimulation monopolar electrode filled with aCSF, placed at about 50–100 μm from the recorded HVC_X_ neuron. Synaptic responses were monitored at different stimulation intensities prior to baseline recording. “Normal” stimulation was defined as a stimulation reliably evoking a synaptic current in the range 100 pA to 1 nA.

Excitatory and inhibitory **synaptic** currents were recorded in the whole-cell voltage clamp mode with the Cs-based patch pipette solution in order to measure the E/I ratio. Only recordings with series resistance below 20 MΩ were included. Excitatory post synaptic currents (EPSCs) and inhibitory post synaptic currents (IPSCs) were recorded at the reversal potential for IPSCs (+10 mV) and EPSCs (−60 mV) in the presence of the NMDA receptor antagonist APV (100 μM), respectively. We also used the Cs-based pipette solution to measure the ratio between N-methyl-D-aspartate-receptor-mediated currents (INMDA) and alpha-amino-3-hydroxy-5-methyl-4-isoxazolepropionic-acid-receptor-mediated currents (IAMPA) in HVC_X_ neurons. We added the GABA_A_ receptor antagonist Picrotoxin (10 μM) to the aCSF for these recordings. I_AMPA_ was recorded at a holding potential of V_h_ = − 60 mV and measured at their peak. I_NMDA_ were recorded in the same cell at Vh =+ 40 mV. I_NMDA_ amplitude was calculated as the mean between 95 and 105 ms after the electric stimulation artifact, to minimize the possible contamination by I_AMPA_.

Access resistance (10–20 MΩ) was monitored throughout the experiment.

#### Intrinsic excitability

Neuronal intrinsic excitability was examined with the potassium gluconate-based pipette solution. After whole-cell current clamp mode was achieved, cells were maintained at −80 mV. Input resistances were monitored by injecting a 150-ms hyperpolarizing current (40pA) to generate a small membrane potential hyperpolarization from the resting membrane potentials. Firing rate represents the average value measured from 1 to 3 cycles (700ms duration at 0.1 Hz, –200 to +250pA range with 50pA or 40pA steps increment, every 12 s).

### Western Blotting in HVC

Samples for western blotting were collected over 2 days between 9am and 12pm from 90-92 dph birds which received either a control (rAAV9/ds-CBh-GFP) or FoxP1 knock-down (pscAAV-GFP-shFoxP1) injection into HVC. Briefly, birds were anesthetized with isoflurane, and their brains were dissected. 230 μm thick sagittal sections were collected in aCSF using a Vibratome VT1200. After confirming the GFP fluorescence under a microscope, HVC was dissected out from these sections. PBS was pipetted out, and the tissue was lysed in RIPA buffer containing protease and phosphatase inhibitors. Protein quantification was done using a Bradford assay, and 50 μg protein/well was used for Western blots. Proteins were run on a 10% SDS-PAGE resolving gel, with 5% stacking gel at 80V until the loading dye front ran off the gel, and were then transferred to an Immuno-Blot PVDF Membrane (Bio-Rad Laboratories) at 250 mA for 2 hr at 4°C. The membrane was dried at room temperature for 1 h, reactivated in methanol, and blocked for 1 hr in 5% milk in TBS. The membrane was cut above 50 kDa, and incubated with primary antibodies in 5% milk in TBS with 0.1% Tween-20 (TBS-T) overnight at 4°C. The following day, it was washed in TBS-T, incubated with secondary antibodies in 5% milk in TBS-T for 1 hr at room temperature, washed in TBS-T again, and imaged with TBS on an Odyssey Infrared Imaging System (LI-COR Biosciences). The following antibodies were used: rabbit α-FOXP1 (Spiteri et al., 2007) (1:5000), mouse α-GAPDH (#MAB374, Millipore, 1:10,000), donkey α-rabbit IgG IRDye 800 (#926-32213, LI-COR Biosciences, 1:20,000), donkey α-mouse IgG IRDye 680 (#926-68072, LI-COR Biosciences, 1:20,000). The images were quantified using the Odyssey Imaging System (LI-COR Biosciences).

### Western Blotting in 293T cells

293T cells were cultured in high-glucose DMEM containing 4 mM L-glutamine, 10% fetal bovine serum (#10437028, Invitrogen), and 1% antibiotic-antimycotic (#15240-062, Invitrogen) at 37°C under 5% CO2. Cells were plated in 6-well plates and transfected with 0.1 pmol/well of overexpression construct (pCMV-FoxP1) and 0.4 pmol/well of hairpin constructs (pscAAV-GFP-shFoxP1 or shCtrl) using FuGENE 6 Transfection Reagent (#E2691, Promega) according to the manufacturer’s instructions. After 24-48h, cells were washed twice with PBS and collected into tubes by scraping in Lysis Buffer (50 mM Tris pH 8.0, 0.1 mM EDTA, 250 mM NaCl, 50 mM NaF, 0.1 mM NaVO4, 10% glycerol, 0.5% IGEPAL CA-630, 1 mM DTT, 0.7 mM PMSF). Samples were incubated for 10 min at 4°C with rotation, and centrifuged at 15,000 rpm for 10 min at 4°C. The supernatant was quantified using a Bradford assay, mixed with loading buffer, incubated at 95°C for 5 min, and stored at −20°C. Western blotting was performed as described for HVC tissue, and imaged on an Odyssey Infrared Imaging System (LI-COR Biosciences).

### Immunohistochemistry

Immunohistochemistry experiments were performed following standard procedures. Briefly, birds were anesthetized with Euthasol (Virbac, TX, USA) and transcardially perfused with PBS, followed by 4% paraformaldehyde in PBS. Free-floating sagittal sections (40 μm) were cut using a cryostat (Leica CM1950, Leica). Sections were first washed in PBS. The tissues were then blocked in 5% normal donkey serum in PBST (0.3% Triton-X in PBS) for 1 hr at RT, incubated with primary antibodies diluted in the blocking buffer (5% donkey serum in PBST), first for 1 hr at RT, and then at 4°C for 48 hrs. Slices were then washed with PBS, and incubated with fluorescent secondary antibodies (diluted in blocking buffer) at RT for 2 hours. After a final PBS wash, sections were mounted onto slides with Fluoromount-G (eBioscience, CA, USA). Composite images were acquired and stitched using an LSM 880 laser-scanning confocal microscope (Carl Zeiss, Germany). Primary antibody: mouse anti-FoxP1 (ab32010, Abcam), 1:1000 concentration. Secondary antibody: donkey anti-mouse, conjugated to AlexaFluor405 (ab175658, Abcam). All image analysis was done using ImageJ, and graphs were prepared in GraphPad Prism7.

### Song analysis

A single day of songs from each FP1-KD behavioural imitation (n=9), FP1-KD social experience (n=8) and control social experience bird (n=10) was selected for analysis when the birds were between 90-100dph. Adult isolate songs were obtained from 8 birds ranging in age from 99-463 dph. Birds were recorded using Sound Analysis Pro 2011 (Tchernichovski et al., 2000).

### Syntax Analysis

From the single day of song for each bird, a random subset of 30 song files were selected for syllable labeling. Syllables were labeled by hand by an expert based on the song spectrograms using a custom MATLAB program. Syllable labels, onset times, and offset times were exported to R for further analysis. Entropy rate was calculated using the *ccber* package in R (Davis et al., 2017) for each bird based on the syllable labels, including a label reflecting the start or end of a song file. Entropy rate reflects an overall measure of the predictability of syllable transitions as a first order Markov chain, calculated according to the formula:

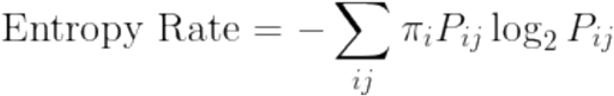

Where *P_ij_* is the probability of transitioning from state *i* to state *j*, and *π_i_* is the stationary distribution of the model for state *i*. A perfectly predictable sequence, where every syllable is always followed by the same syllable will have an entropy score of 0. A maximally entropic syllable sequence, where there is an equal probability that a given syllable transitions to any of the possible syllables (or a file end), will have an entropy rate of log_2_*K*, where *K* is the number of different states, or in this case syllables plus an additional state representing the end of a file.

Syllable transition probabilities used to determine arrow thickness in Figure 2d, h, l & p were calculated using the *markovchain* package in R (Spedicato & Signorelli, 2014). Each syllable, as well as file boundaries and gaps lasting longer than 100ms were considered ‘inter-bout gap’ states in the first order Markov chain. Each pairwise transition between states (syllables or inter-bout gaps) is tallied to determine the probability of each of the possible states occurring following a given current state in order to generate a two-dimensional transition probability matrix. Arrow thicknesses in Figure 2d, h, I & p are directly proportional to the probability that the origin syllable would transition to the syllable designated by the arrow.

Syntax raster plots shown in Figure 2 c, g, k & o were created using a custom R code. Syllable labels are first arranged into bouts. Bouts are considered strings of at least two syllables which are not separated by a gap longer than 100ms or a file boundary. All bouts are then arranged according to user-specified primary and secondary alignment syllables. These alignment syllables are chosen to maximize the overall alignment of all bouts in the final raster plot. Each bout is shifted along the x axis such that the first occurrence of the primary alignment syllable occupies the 0 position across all bouts, with the order of syllables maintained within each bout. If a bout does not contain the primary alignment syllable, it is shifted such that the first secondary alignment syllable occupies the 0 position, and the order of syllables within the bout is maintained. Bouts which contain neither the primary nor secondary alignment syllables are plotted above the others at an offset, such that the last syllable in that bout occupies the x position −1. Once aligned, the bouts are ordered along the y axis alphabetically based on the syllable labels following the alignment syllable. The result is a representation of the sequence of syllables across all labeled bouts from a single bird arranged to maximize the emergence of patterns and dominant sequences.

### Similarity Scoring

The lack of stereotyped motif structure in the isolate tutored shFoxP1 birds made it difficult to use standard automated song similarity scoring programs. Instead, similarity between spectrograms was evaluated by 11 human experts. Each was presented with a total of 126 spectrogram pairs, and were instructed to rate the similarity of the two spectrograms on a scale of 1 (not similar) to 10 (very similar), following a 5 comparison training set. The spectrograms used were all generated using Sound Analysis Pro 2011 (Tchernichovski et al., 2000) and all reflected a duration of approximately 2.5 seconds. All spectrograms began at the beginning of a song bout, and with the exception of songs which lacked a typical motif structure, included at least one full song motif. The order of comparisons within the test set was randomized for each participant, and included 4 comparisons between tutee and tutor for each tutee in the FP1-KD behavioural imitation, FP1-KD social experience and control social experience groups (4 x 27 birds), 8 comparisons between a tutor and itself to ensure that scorers were using the full range of the scale, and 10 duplicated tutor-tutee pairs to ensure that the scorers were internally consistent.

No individual scorer differed from the mean score by more than an average of 2 standard deviations, none differed by more than an average of 2 points on the duplicated comparisons, and all but one made use of the full 10 point scale (they gave at least one comparison a score of 0 and at least one a score of 10). One scorer never gave any comparison a perfect 10/10 score, so their scores were rescaled such that they spanned the full 1-10 range.

A % similarity score was calculated for each bird by taking the mean similarity rating for all 4 comparisons to the tutor across all scorers.

The full scoring set is available at https://forms.gle/9TDu1fwGGYXWKhgB6.

### Statistical Analysis

All data were tested for normality using the Shapiro-Wilk Test. Consequently, parametrical or non-parametric statistical tests were used as appropriate: t-test (unpaired or paired, as indicated), Mann-Whitney (unpaired) or Wilcoxon matched-pairs signed rank test (paired) tests were used where appropriate. One-way Anova or Kruskal-Wallis test was performed when comparisons were made across more than two conditions. Two-way Anova (post-hoc Sidak) was used to test differences between two or more groups across different conditions. Statistical significance refers to \**P* < 0.05, ** *P* < 0.01, *** *P* < 0.001. Statistical details for all experiments are included in the corresponding figure legends.

**Extended Data Figure 1:**
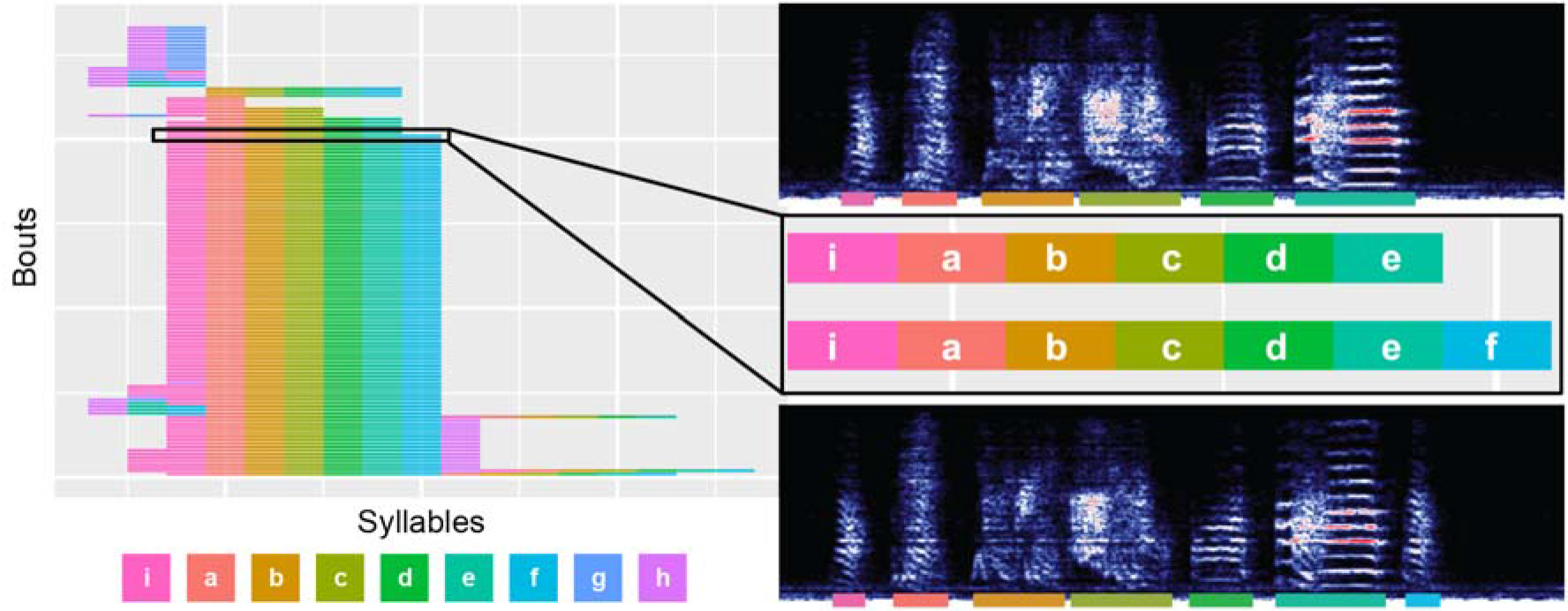
Interpreting a syntax raster plot. Schematic, featuring song data from the non-isolate shFoxP1 bird featured in figure 2**b-d**, which illustrates the relationship between raw spectrograms and syntax raster plot visualizations used in figures 1 and 2. The syntax raster plot represents 30 randomly selected song files from a single bird. These files are segmented into syllables by amplitude thresholding, then syllables are labelled by hand. The song files are then separated into song bouts, defined as more than two syllables produced consecutively with a gap shorter than 100ms between them. These bouts are then sorted according to the syllables they contain so that identical bouts appear together. Each song bout is then represented as a row of blocks whose colour reflects the syllable label at each position within the bout. Birds with very stereotyped syllable order, such as the one in this figure, will have raster plots with a consistent vertical stripe pattern, indicating that the same syllable is consistently produced in the same position.

**Extended Data Figure 2:**
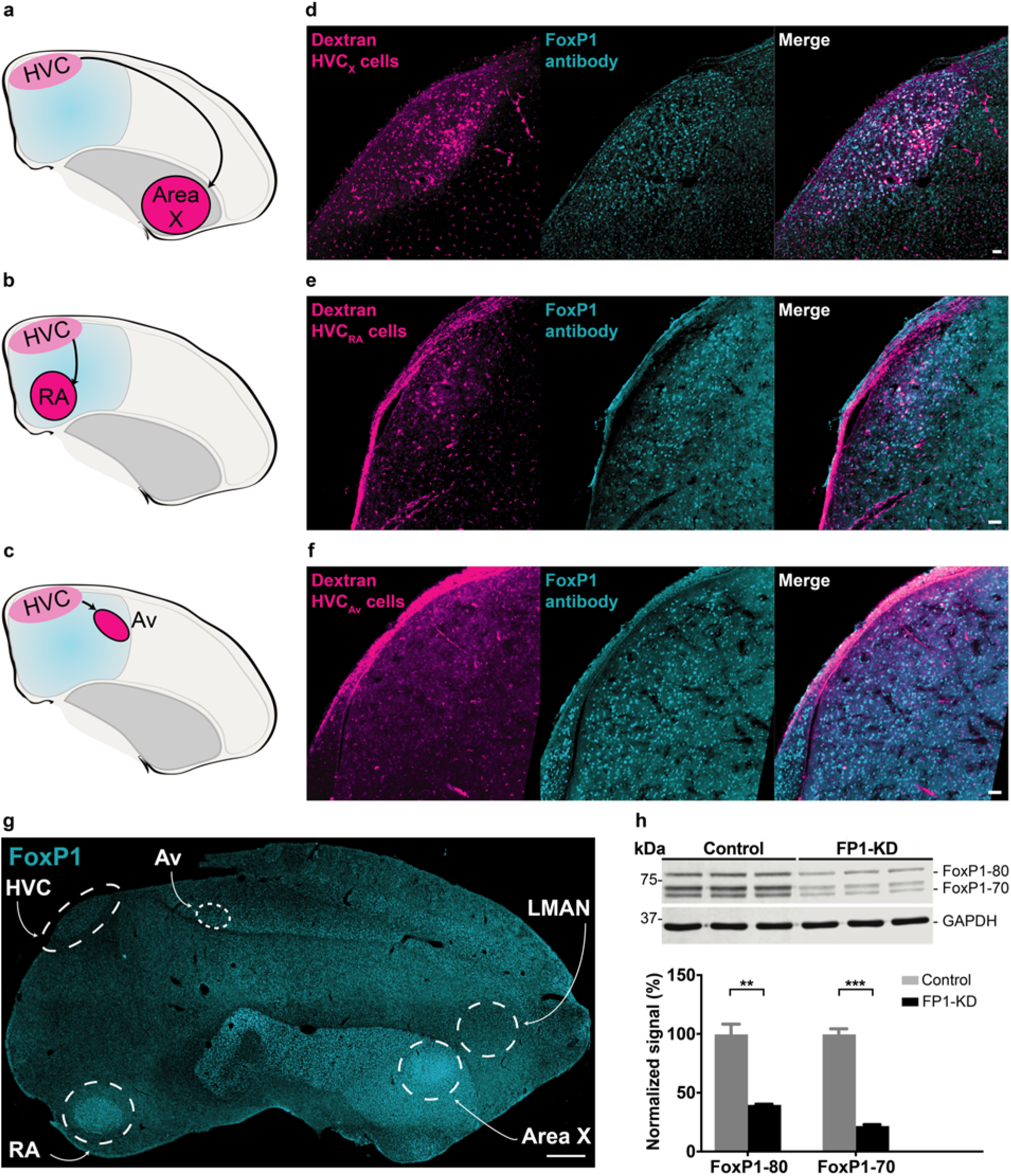
FoxP1 expression differs between HVC subtypes, various song nuclei, and in cells overexpressing zfFoxP1. **a-f,** FoxP1 labelling in various subtypes of HVC projection neurons: HVC_X_ (**a,d**), HVC_RA_ (**b,e**), and HVCAv (**c,f**). **a-c,** Schematic of Dextran injections for retrograde labelling of the various HVC neuronal subtypes; **d-f,** Examples of sagittal sections through HVC showing retrograde labelling with Dextran-594 (magenta, left), immunolabelling with FoxP1 (cyan, middle), and a composite image showing coexpression of FoxP1 in the respective HVC neurons (merge, right). Scale bars, 50 μm. **g,** FoxP1 staining in a sagittal brain slice: FoxP1 is prominently present in Area X, HVC, RA and the mesopallium (including Av), and absent in LMAN (as in Mendoza et al., 2015). Scale bar, 500 μm. **h,** (top) Western blot of lysates from 293T cells overexpressing FoxP1, and either control (n = 3) or shFoxP1 (n = 3) AAV constructs. At least two isoforms of FoxP1 were observed around 80 kDa and 70 kDa, with GAPDH as loading control. (bottom) Quantification of top blot. FoxP1 protein signals were normalized to GAPDH, averaged for each condition, and normalized to Controls. Histograms represent average±SEM% of the normalized signal. Cells transduced with pscAAV-GFP-shFoxP1 (FP1-KD) showed significantly reduced FoxP1 expression than those with the control virus. (<u>FoxP1-80</u>: Control: 100±8.9%, n=3, vs FP1-KD: 40.0±0.7%, n=3, Student’s t-test, *P*= 0.0025; FoxP1-70: Control: 100±4.8%, n=3, FP1-KD: 22.0±1.3%, n=3, Student’s t-test, *P* < 0.001).

**Extended Data Figure 3:**
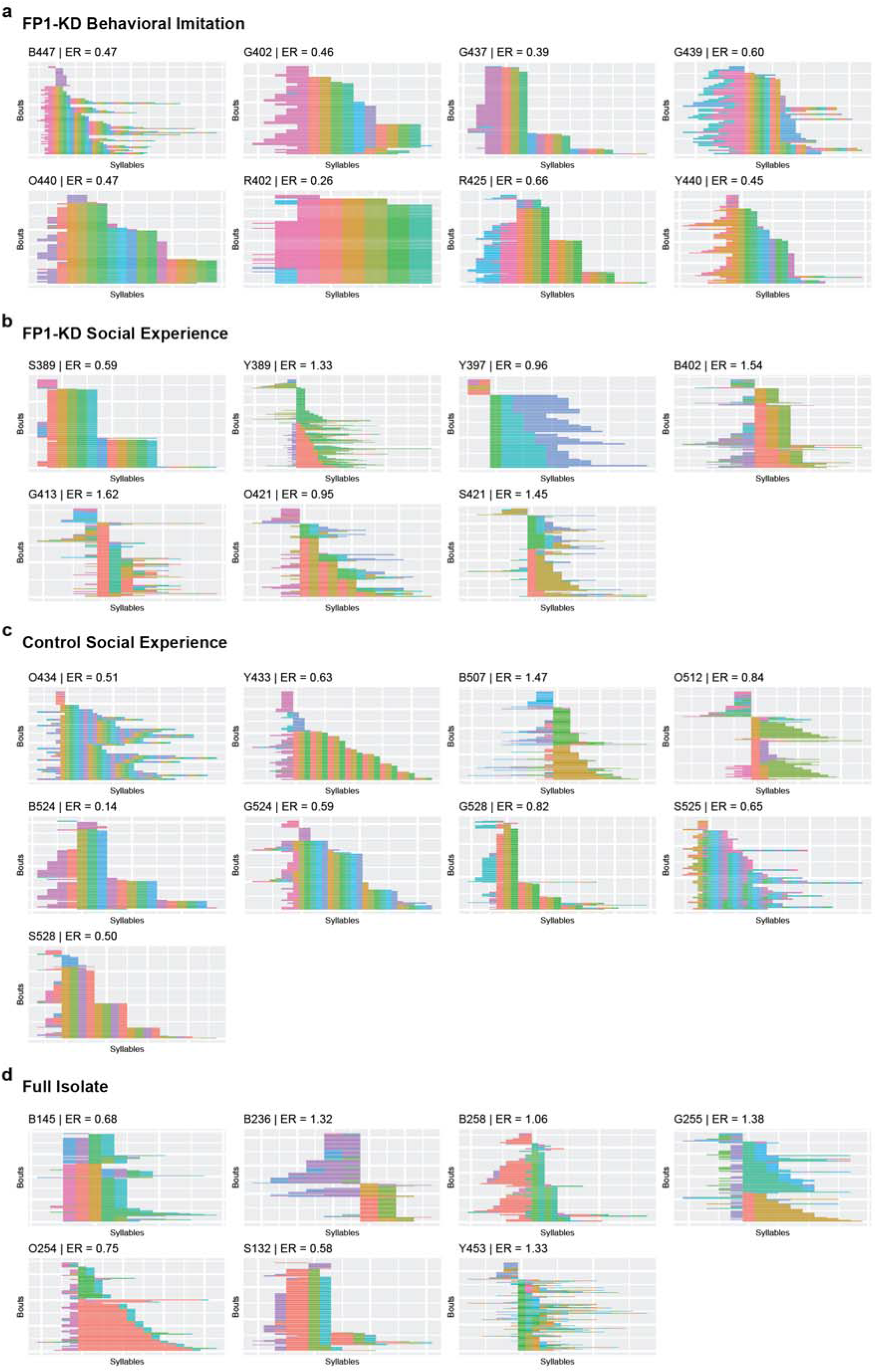
HVC FoxP1 knock-down in isolate birds impairs subsequent song sequence stereotypy. Syntax raster plots from each bird included in song syntax analysis. ER refers to the first order Markov entropy rate score for that individual bird.

**Extended Data Figure 4:**
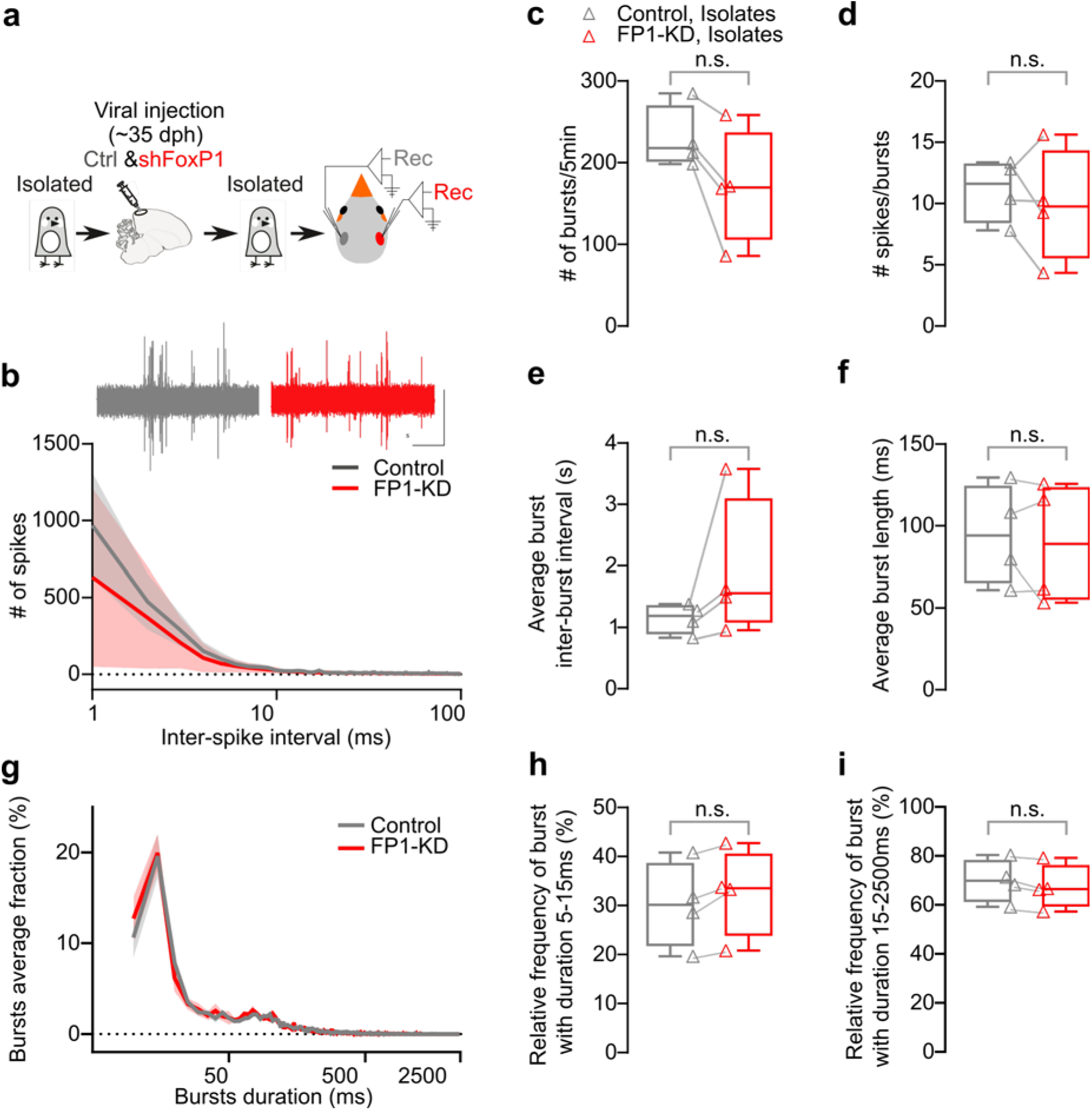
Unilateral HVC FoxP1 knock-down does not modify in-vivo extracellular recorded spiking and bursting pattern in isolated birds. **a,** Schematic representing the experimental timeline. Birds received AAV injections in HVC to knock-down FoxP1 in one hemisphere, and control virus in the other hemisphere (pseudo-randomized). The birds were kept in isolation from male tutors, and ~10 days later, HVC extracellular activity in both hemispheres (n = 4 birds, 3-5 recordings/hemisphere) was recorded. **b,** (top) Sample traces (scalebar 0.5V, 1s) and (bottom) average inter-spike interval distribution (bin 1ms, 1-100ms, logarithmic scale, 300s/recording) (Control vs FP1-KD hemispheres, Two-Way Anova F_1,6_ = 0.7424, *P* > 0.05). **c,** Box- and scatter plots of the total number of bursts (Control hemispheres: 229.6±19.1 vs FP1-KD hemispheres: 170.8±35.2, Wilcoxon matched-pairs signed rank test, *P* > 0.05). **d,** Box- and scatter plots of the average number of spikes in a burst (Control hemispheres: 11.1±1.3 vs FP1-KD hemispheres: 9.9±2.3, Wilcoxon matched-pairs signed rank test, *P* > 0.05). **e,** Box- and scatter plots of the average inter-burst interval (Control hemispheres: 1.1±0.1 vs FP1-KD hemispheres: 1.9±0.6, Wilcoxon matched-pairs signed rank test, *P* > 0.05). **f,** Box- and scatter plots of the average burst length (Control hemispheres: 94.6±15.2 vs FP1-KD hemispheres: 89.2±18.5, Wilcoxon matched-pairs signed rank test, *P* > 0.05). **g,** Plot representing the relative distribution of bursts duration, normalized for each recording (5ms duration bins, 5-2500ms, logarithmic scale; Control hemispheres vs FP1-KD hemispheres: Two-Way Anova, Interaction F_498,2988_ = 0.05189, *P* > 0.05). **h,** Box- and scatter plots of the average relative prevalence of bursts with duration comprised between 5 and 15ms (Control hemispheres: 30.2±4.4 vs FP1-KD hemispheres: 32.6±4.5, Wilcoxon matched-pairs signed rank test, *P* > 0.05). **i,** Box- and scatter plots of the average relative prevalence of bursts with duration comprised between 15 and 2500ms (Control hemispheres: 69.8±4.4 vs FP1-KD hemispheres: 67.4±4.5, Wilcoxon matched-pairs signed rank test, *P* > 0.05).

**Extended Data Figure 5:**
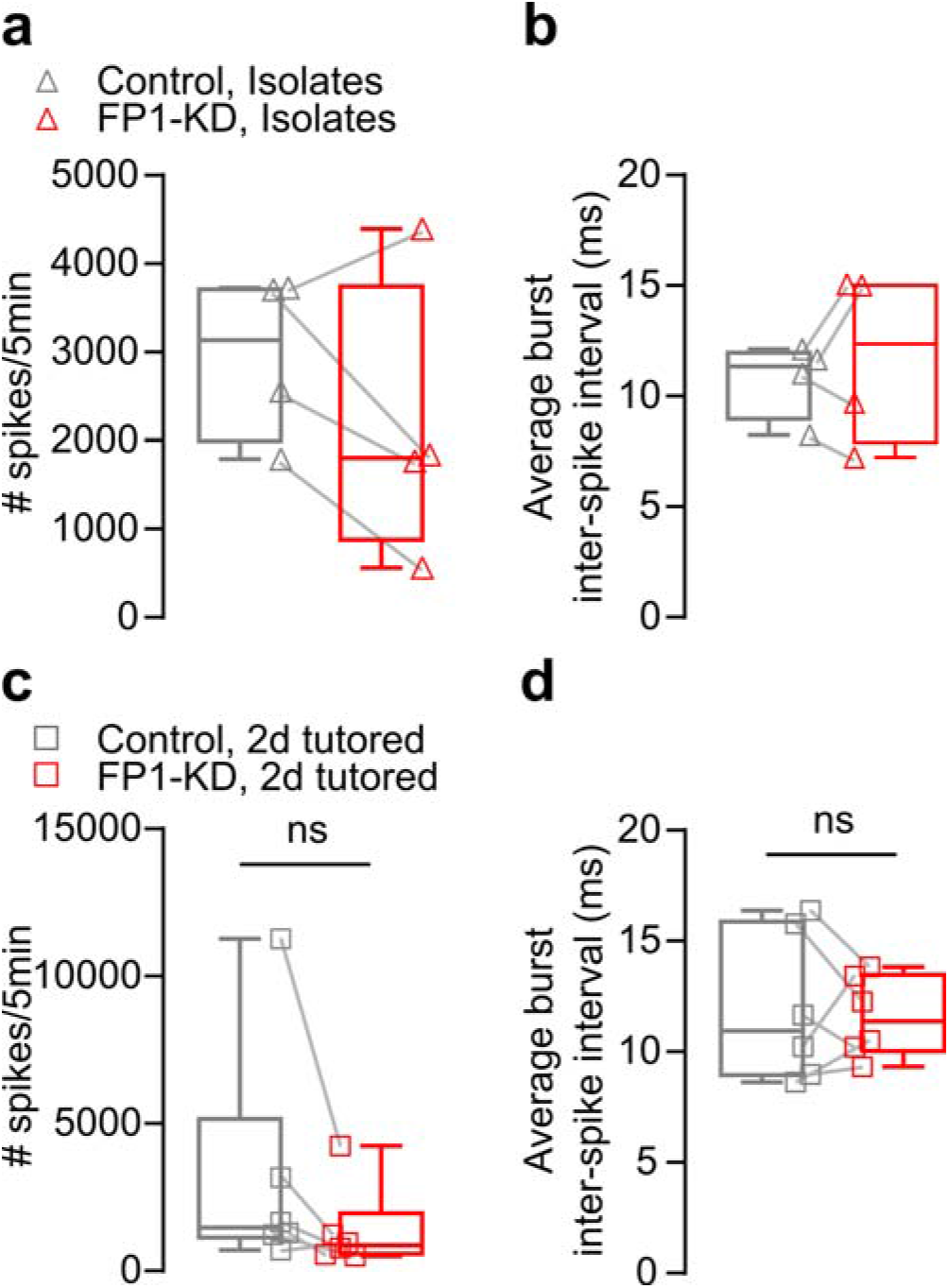
Spiking properties in unilaterally-injected FP1-KD birds remain unchanged, irrespective of previous social experience. **a,**Box- and scatter plots of the average number of spikes recorded in isolates FP1-KDshFoxP1 and Control hemispheres (Control hemispheres: 2946±472 vs FP1-KD hemispheres: 2141±806, Wilcoxon matched-pairs signed rank test, *P* > 0.05). **b,** Box- and scatter plots of the average inter-spike interval in bursting events recorded in isolates FP1-KD and Control hemispheres (Control hemispheres: 10.8±0.9 vs FP1-KD hemispheres: 11.8±2.0, Wilcoxon matched-pairs signed rank test, *P* > 0.05). **c,** Box- and scatter plots of the average number of spikes recorded in 2d-tutored FP1-KD and Control hemispheres (Control hemispheres: 3222±1646 vs FP1-KD hemispheres: 1379±584, Wilcoxon matched-pairs signed rank test, *P* > 0.05). **d,** Box- and scatter plots of the average inter-spike interval in bursting events recorded in 2d-tutored FP1-KD and Control hemispheres (Control hemispheres: 11.9±1.4 vs FP1-KD hemispheres: 11.6±0.8, Wilcoxon matched-pairs signed rank test, *P* > 0.05).

